# Graph neural networks best guide phenotypic virtual screening on cancer cell lines

**DOI:** 10.1101/2024.06.26.600790

**Authors:** Sachin Vishwakarma, Saiveth Hernandez-Hernandez, Pedro J. Ballester

## Abstract

Artificial intelligence is increasingly driving early drug design, offering novel approaches to virtual screening. Phenotypic virtual screening (PVS) aims to predict how cancer cell lines respond to different compounds by focusing on observable characteristics rather than specific molecular targets. Some studies have suggested that deep learning may not be the best approach for PVS. However, these studies are limited by the small number of tested molecules as well as not employing suitable performance metrics and dissimilar-molecules splits better mimicking the challenging chemical diversity of real-world screening libraries. Here we prepared 60 datasets, each containing approximately 30,000 to 50000 molecules tested for their growth inhibitory activities on one of the NCI-60 cancer cell lines. We evaluated the performance of five machine learning algorithms for PVS on these 60 problem instances. To provide a comprehensive evaluation, we employed two model validation types: the random split and the dissimilar-molecules split. The models were primarily evaluated using hit rate, a more suitable metric in VS contexts. The results show that all models are more challenged by test molecules that are substantially different from those in the training data. In both validation types, the D-MPNN algorithm, a graph-based deep neural network, was found to be the most suitable for building predictive models for this PVS problem.

## 1 Introduction

Recent years have witnessed strong advancements in computational drug discovery methodologies. Targeted Drug Discovery (TDD), which aims at identifying molecules that interact with a specific molecular target associated with the considered disease [1], [2], [3], has been the predominant approach. Gradually, however, Phenotypic Drug Discovery (PDD) approaches have gained traction [4], [5], [6]. Distinct from TDD, PDD does not depend solely on the molecular understanding of diseases; rather, it focuses on observable phenotypic changes, offering a more expansive approach that is not confined to known targets [7], [8]. This has the advantage of enabling the exploration of therapeutic agents through mechanisms of action that are not yet known, potentially leading to innovative treatments. Moreover, PDD acknowledges that molecules effective at targeting specific processes in isolation may also perform effectively within a broader cellular environment [9], [10]. This is important as many molecules with potent activity for a target are later found to lack whole-cell activity for a range of reasons, e.g., the target not being co-located with the molecule in the cell [9].

High-Throughput Screening (HTS) techniques have been fundamental to both TDD and PDD, enabling the assessment of libraries with up to a few million compounds [11], [12]. However, as the scale of chemical libraries grows to giga-scale proportions, HTS is no longer an option to screen them. This challenge is compounded by the increasing complexity of cancer as a multi-genic disease, which requires the exploration of vast, uncharted chemical spaces to identify effective therapeutics.

In this context, Artificial Intelligence (AI) presents a tremendous opportunity to transform the landscape of early drug discovery [13], [14]. AI’s capacity to leverage existing data from HTS to predictively model and explore vast chemical libraries is transforming drug screening [15]. Graph Neural Networks (GNNs), in particular, have shown promise owing to their ability to model complex, irregular data structures inherent in molecular chemistry [16]. Yet, despite AI’s advancements, there remain significant limitations in its performance, especially when tasked with predicting responses from chemically dissimilar molecules [17], [18],[19]. These limitations highlight a critical gap in current AI methodologies, which often fail to generalize beyond the chemical entities included in their training sets (unseen molecules). This is particularly the case when predicting the properties of novel active compounds (i.e. not only unseen but also dissimilar to the known actives of the considered target).

The efficacy of AI models in the activity of molecule-cell line pairs has been evaluated in various contexts using comprehensive databases such as GDSC, CCLE, and NCI-60 [17], [18], [19], [20], [21], [22], [23], [24], [25], [26], [27]. These evaluations mostly focus on predicting the responses of unseen cell lines to a given drug, e.g. [28], [29], [30], [31], [32], for precision oncology purposes. A few other evaluate such phenotypic drug response models on unseen drugs [18], [19], [20], [31], [33], and leave-drug-out scenarios [20], [28], [31], [33], [34] for a given cell line. However, the evaluation of AI models on test sets with chemically dissimilar molecules remain significantly underexplored [18]. Another shortcoming is that practically all these studies do not evaluate the developed models for virtual screening (here discriminating between active and inactive molecules on the considered cell line). This is partly due to the most popular resources only having a few hundred drugs being tested on each cell line (e.g. GDSC, CCLE).

As a consequence of these shortcomings, we still do not know which are the best AI models to guide virtual screening of gigascale libraries against cancer cell lines. Indeed, Machine Learning (ML) methods for tabular datasets like Random Forests (RF) and XGBoost (XGB) often yield better performance with smaller datasets due to their efficiency in handling limited data [18], [26], [35], highlights the need for more comprehensive studies on how different AI models, particularly GNNs, perform when applied to test sets with more realistic chemical diversity [34], [36].

To answer these research questions, we will assemble 60 datasets, each with diverse molecules tested on one of the cell lines in the NCI-60 panel [37]. Then, we will investigate how each algorithm performs across these 60 instances of the same problem, highlighting differences in algorithmic efficiency and predictive accuracy in a controlled yet varied set of conditions. The NCI-60 tumour cell line panel, with its extensive profiling of over 130,000 compounds, underscores the critical role of advanced data models in refining drug sensitivity predictions [31], [38], [39]. By utilizing such comprehensive datasets, this study aims to rigorously assess the performance of supervised learning algorithms in predicting the biological activity of chemically dissimilar molecules against cancer cell lines, an area yet to be fully explored by current AI-driven methodologies [28]. This will be carried out with appropriate evaluation metrics, such as the Hit Rate [15], [18], [40], which are more suited for virtual screening than traditional ROC-AUC measures [21], [27], [28], [34].

## 2 Materials and Methods

### 2.1 NCI-60 dataset

The dataset employed in this research was obtained from the NCI-60 database, which is publicly available at https://wiki.nci.nih.gov/display/NCIDTPdata/NCI-60+Growth+Inhibition+Data. This comprehensive dataset includes growth inhibition information for a wide array of 159 cell lines tested against 53,215 molecules (distinct NSC IDs). In total, the dataset encompasses 3,054,004 measurements of pGI_50_, which quantifies the logarithmic concentration of a compound necessary to achieve a 50% reduction in tumour growth. To enhance the reliability of our analysis, we excluded 41,113 pGI_50_ values below the threshold of 4, considering them less reliable due to extrapolation from higher concentration ranges. In cases of multiple pGI_50_ values for the same NSC-Cell line pair, we computed the average, recognizing that potent molecules often undergo retesting across various concentration ranges, leading to multiple pGI_50_ recordings.

The processed dataset comprises 60 cell lines, shown in Figure 1. Molecular representations were converted from SDF to SMILES format utilizing the Open Babel library [41]. The chemical structures were then standardized using the Molvs package (https://molvs.readthedocs.io/en/latest/). This standardization procedure involved sanitization, hydrogen removal, metal disconnection, application of normalization rules, acid re-ionization, and stereochemistry recalculations. Following this, 1,137 NSCs were omitted from the dataset due to the unavailability of chemical descriptors.

**Figure 1.**
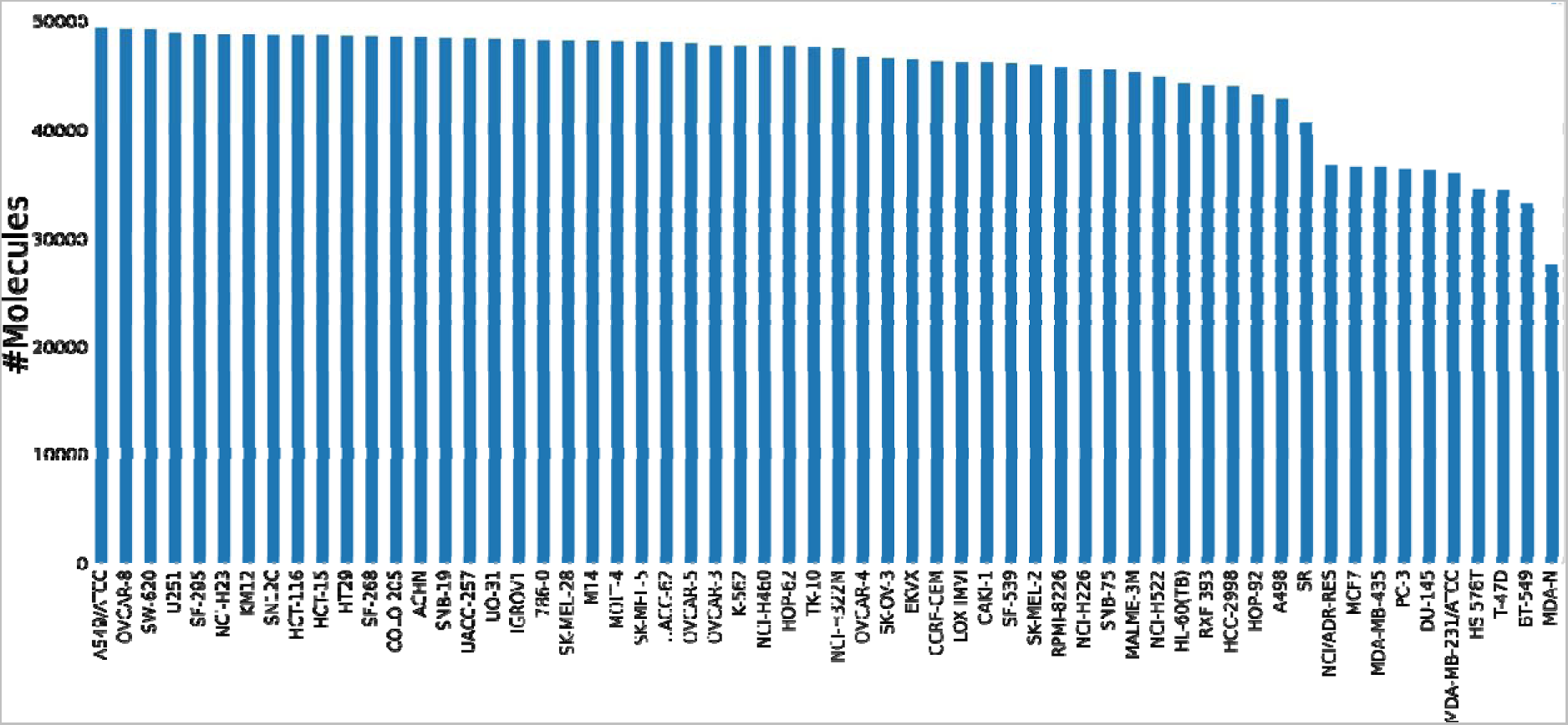
Distribution of molecules tested across NCI-60 cell lines. The x-axis represents the cell lines, while the y-axis indicates the number of molecules tested on each cell line. The vertical bars show the variation in the number of molecules tested across different cell lines, providing insights into the diversity of experimental conditions and the comprehensiveness of the study. The data represents the distribution of the number of pGI_50_ measurements, one per cell line-NSC pair, across the 60 cell lines.

Our analysis primarily focused on 60 extensively profiled cell lines, as documented in previous cancer research studies [18], [19], [26], [31], [42]. The NCI-60 has profiled data spanning nine cancer types, including Leukaemia (LE), Melanoma (ME), Non-small-cell Lung (LC), Colon (CO), Central Nervous System (CNS), Ovarian (OV), Renal (RE), Prostate (PR), and Breast (BR). The refined dataset encompasses 2,707,434 unique NSC-Cell line pairs, incorporating data for 60 cell lines and 50,555 molecules (totalling 50,846 unique NSC IDs). A discrepancy in the count of unique canonical SMILES (50,156) relative to unique NSC IDs highlights instances where different NSC IDs were assigned identical SMILES representations, underscoring the importance of canonical SMILES as a unique molecular identifier.

In our study, we opted to use Morgan circular fingerprints [43] to represent the molecular features of compounds, leveraging the RDKit library (https://www.rdkit.org/) for their generation in bit vector format. The choice of Morgan fingerprints was informed by their demonstrated efficacy in enhancing model performance across similar research endeavours [19], [31], [44]. These fingerprints effectively capture the presence or absence of specific substructures within a molecule, offering a comprehensive molecular representation. The configuration of the fingerprint’s radius and bit size are pivotal in generating meaningful Morgan circular fingerprints. In our analysis, we utilized a bit size of 256 with a radius of 2, aligning with configurations that have previously yielded optimal results in the literature [19], [31].

Beyond structural features, we also quantified several physicochemical properties of the compounds to enrich our feature set. These properties encompassed: Total Polar Surface Area (TPSA) [45], Molecular Weight (MW) [46], LogP (Partition coefficient) [47], Number of Aliphatic Rings, Number of Aromatic Rings [48], Number of Hydrogen Bond Acceptors, Number of Hydrogen Bond Donors [49]. The inclusion of these physicochemical descriptors alongside Morgan fingerprints provides a multidimensional representation of each compound, contributing to the robustness and predictive capability of our modelling approach.

### 2.2 Machine learning

#### 2.1.1 Linear Regression

We used linear regression (LR) to establish a baseline for comparing model performance. As implemented in the scikit-learn library [50], the LR model predicts an output (*y*) by applying a linear combination of input variables (Χ). This relationship is mathematically represented as:

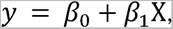

where β_0_ is the intercept (or bias coefficient), and β_1_ is the weight assigned to each input feature. For scenarios encompassing multiple input variables, the model extends beyond a simple line to form a hyperplane. This multidimensional approach allows for a more nuanced representation of the data, accommodating complex relationships between multiple inputs and the predicted output. The configuration of the equation, particularly the values and interactions of its coefficients (β_0_ and β_1_ etc.) visually conveys the model’s interpretation of the underlying problem.

#### 2.1.2 Random Forest

RF is an ensemble learning method that improves prediction accuracy and controls overfitting by creating a forest of decision trees [51], [52]. RF employs the bagging (Bootstrap aggregating) technique [53] with a distinct approach of using only a subset of features for splitting each node in the decision trees. This method ensures that the trees within the forest are uncorrelated, enhancing the ensemble’s prediction quality by reducing variance without significantly increasing bias. RF aggregates predictions from each tree to minimize prediction error and mitigate the impact of outliers and noise.

In our study, we optimized RF through the tuning of two hyperparameters: *n_estimators,* the number of trees in the forest, and *max_features*, the number of features considered for splitting at each node. We adopted the bootstrap sampling technique and employed the Mean Squared Error (MSE) as the scoring function to assess model performance on the test set. We initially configured the *n_estimators* parameter to 500 and set *max_features* to 0.33 based on recommendations [51]. The *max_depth* parameter was left unrestricted. This setup served as a foundation for further hyperparameter tuning to identify the optimal hyperparameter range that enhances model performance. Table 1 shows the examined range of hyperparameters for the RF algorithm.

**Table 1.** Values tested/searched for each hyperparameter of the ML algorithms.

|  |  |  |  |
| --- | --- | --- | --- |
| Algorithm | Hyperparameters | Values tested | Tuning method |
| RF | N_estimators | 250, 500, 750, 1000 | Grid search |
|  | Max_features | 0.2, 0.3, 0.4, 0.6, 0.8, 0.9 |  |
| XGB | N_estimators | 250, 500, 750, 1000 | Grid search |
|  | Max_depth | 5,6,7,8,9,10 |  |
|  | Cosample_bytree | 0.3, 0.4, 0.6, 0.8, 0.9 |  |
| DNN | Learning rate | 0.0001, 0.001, 0.01, 0.1 | Grid search |
|  | Weight decay | 0, 1e-6, 1e-5, 1e-4, 1e-3, 1e-2 |  |
|  | Drop out | 0, 0.2, 0.4, 0.6, 0.8 |  |
| D-MPNN | Hidden size | low=300, high=2400 | Bayesian optimization |
|  | Depth | low=2, high=6 |  |
|  | Dropout | low=0.0, high=0.4 |  |
|  | Ffn_num_layers | low=1, high=3 |  |

#### 2.1.3 Extreme Gradient Boosting

XGB stands out as a sophisticated ensemble learning algorithm, renowned for its efficiency and effectiveness [54]. XGB operates on the principle of boosting, which seeks to iteratively minimize prediction errors using a gradient descent optimization algorithm. Unlike traditional models that train in isolation, XGB enhances model accuracy sequentially, with each new tree aiming to correct the residual errors left by its predecessors. This iterative correction continues until no significant improvement is observed or a pre-determined number of iterations is reached. Distinguishing itself from RF, which averages predictions from independently trained models, XGB focuses on sequential improvement, making each tree dependent on the corrections from the trees before it.

XGB’s performance relies on key hyperparameters, which include *max_depth,* the maximum depth of any decision tree in the ensemble; *n_estimators,* the total number of trees in the ensemble; *colsample_bytree, t*he fraction of features used to train each tree; and *eta,* the learning rate that controls each tree’s contribution to the final outcome. Previous studies have determined initial values for each of these hyperparameters, particularly setting eta to 0.5 [55]. These default values are utilized in our current study to identify the optimal combination for our specific task. Details of the hyperparameters explored can be found in Table 1.

#### 2.1.4 Deep Neural Network

Deep neural networks (DNNs) are more precisely deep multi-layer perceptrons, celebrated for their ability to intricately model complex relationships in high-dimensional data [56]. They significantly enhance our comprehension and interpretation of data through sophisticated, layered computational structures. In this study, we employed the Keras framework with TensorFlow backend, as outlined at (https://keras.io/), to develop a DNN tailored for regression analysis. Our model architecture is fully connected and comprises an input layer, multiple hidden layers, and an output layer, each integral to the network’s predictive.

The architecture of the DNN is the following:

- *Input Layer.* This layer is dimensioned to match the number of input features, ensuring that each feature is adequately represented.
- *Output Layer.* Consisting of a single neuron, this layer is designed for regression, producing a continuous output value.
- *Hidden Layers.* The network includes several hidden layers, which are instrumental in learning the data’s complex patterns. The ‘width’ of a layer refers to its number of neurons, and the network ‘depth’ to its number of layers. Optimization of neuronal weights across these layers is accomplished through the Adam optimizer [57], a decision driven by its effectiveness in gradient descent optimization.

The considered the DNN settings are:

- *Weight Initialization* [58]. Implemented to prevent the vanishing or exploding gradient problem, ensuring efficient forward and backward propagation.
- *Optimization Algorithm.* The core of the training process involves iterative weight adjustments using the Adam optimizer [57], aimed at minimizing the loss function.
- *Activation Functions.* Non-linear activation functions, specifically ReLU [59], [60], for the hidden layers, are employed to introduce non-linearity, enabling the model to capture complex relationships within the data.

Strategies to counteract overfitting include:

- *Dropout* [61]. Randomly omitting neurons during training to encourage more generalized learning.
- *Regularization* [62]. Applying weight decay through L1 regularization to reduce the model’s complexity by penalizing large weights, thereby promoting simpler models.

Our approach to hyperparameter tuning involved a grid search strategy to systematically explore the effects of various parameters, including learning rate, weight decay, dropout rates, and activation functions, on model performance. This process was informed by recent literature [63], [64], guiding the selection of ReLU activation for hidden layers and a linear activation for the output, alongside the implementation of batch normalization [65] and the Adam optimizer to enhance the learning process.

The final model configuration, determined through empirical hyperparameter tuning, consists of three hidden layers with respective neuron counts of 512, 256, and 64. Training was conducted over 100 epochs with a batch size of 100, using the MSE as the loss function. This configuration was selected to optimize the model’s ability to learn from the training data and accurately predict outcomes on unseen data. Table 1 details the range of hyperparameters evaluated.

The optimized hyperparameter set, comprising a learning rate of 0.01, a dropout rate of 0.6, and no weight decay, was chosen based on its performance in enhancing the model’s prediction accuracy and generalization capability. The neuron configuration for the hidden layers was adjusted to 2048, 1024, and 512 to maximize the model’s effectiveness in complex pattern recognition and prediction.

#### 2.1.5 Directed-Message Passing Neural Network

The Directed-Message Passing Neural Network (D-MPNN) [66], [67] is a graph-based Neural Network (NN) architecture adept at learning molecular representations from molecular graphs derived from SMILES (Simplified Molecular Input Line Entry System) inputs [16]. By translating the SMILES into graphical data, D-MPNN incorporates detailed atom and bond information—such as atom type, bond quantity, charge, chirality, and more—that are largely one-hot encoded (Table 2) except for atomic mass which is treated as a scaled real number for consistency across all molecules.

**Table 2.** Atom and Bond features included in the D-MPNN algorithm.

| **Table 2. Atom and Bond features included in the D-MPNN algorithm.** |  |  |  |
| --- | --- | --- | --- |
| **Category** | **Features** | **Description** | **Number of features** |
| Atom features | atom type | Type of atom (C, O, N), by atomic number | 100 |
|  | # bonds | Number of bonds | 6 |
|  | Formal charge | Electronic charge assigned to an atom | 5 |
|  | Chirality | Unspecified, tetrahedral CW/CCW, or other | 4 |
|  | # Hs | Number of bonded hydrogens | 5 |
|  | Hybridisation | sp, sp<sup>2</sup>, sp<sup>3</sup>, sp<sup>3</sup>d or sp<sup>3</sup>d<sup>2</sup> | 5 |
|  | aromaticity | Part of aromatic system or not | 1 |
|  | Atomic mass | Mass of the atom, divide by 100 | 1 |
| Bond features | bond type | Single, double, triple, or aromatic | 4 |
|  | conjugated | Whether the bond is conjugated | 1 |
|  | In-ring | Whether the bond is part of a ring | 1 |
|  | stereo | None, any, E/Z or cis/trans | 6 |

The process, depicted in Figure 2, begins by encoding node descriptors within an adjacency matrix to demonstrate atom connections, where a ‘1’ signifies a direct bond. The D-MPNN operates through two primary phases, message passing and readout phases. During the message passing phase, atom and bond features are aggregated to create a detailed molecular representation. This involves collecting information from neighbouring atoms and bonds, with the depth of message passing—up to three bond lengths in this study—being a tuneable parameter. The readout phase constructs a feature vector for the entire molecular graph by applying a readout function that aggregates the updated atom features, leading to the final molecular embedding used for predictive modelling.

**Figure 2.**
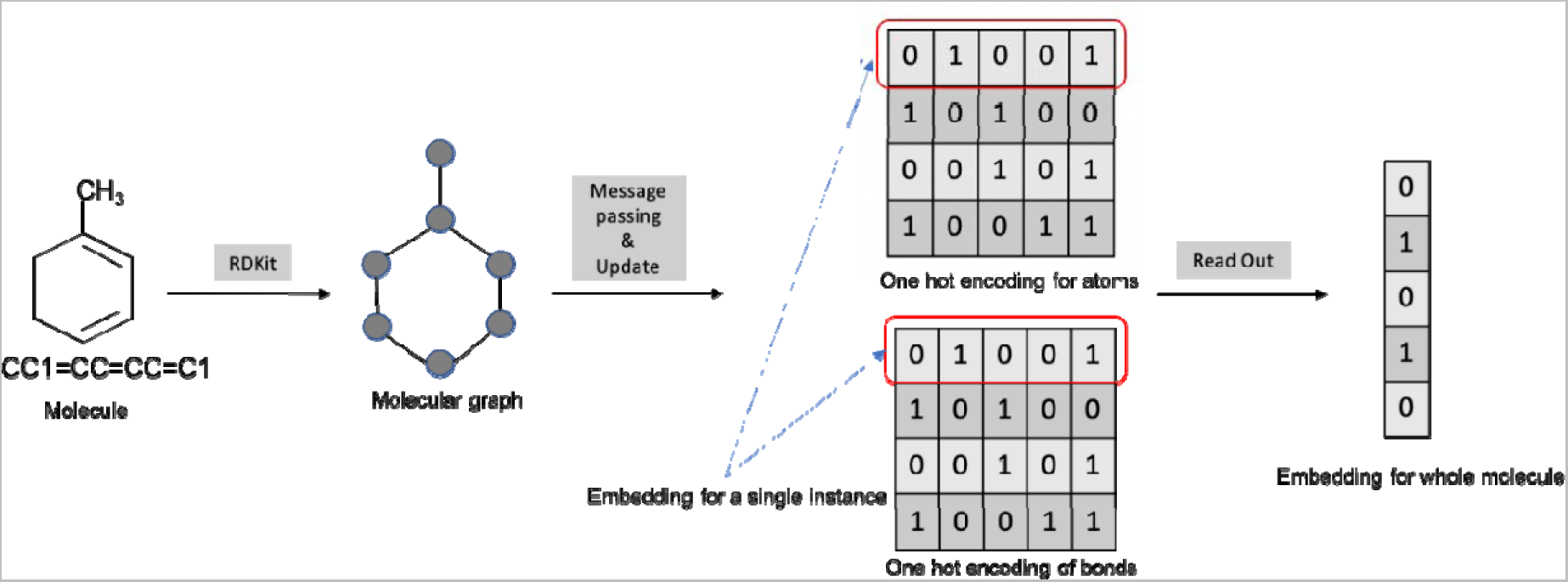
Sequence of operations executed by the D-MPNN algorithm to extract features from chemical compounds. Beginning with a SMILES string, the algorithm employs the RDKit library to construct a 3D molecular structure. This structure enables the algorithm’s message passing and update function to assimilate the attributes of adjacent atoms and bonds. The culmination of this process is the conversion of the assimilated features into a user-defined fixed-length vector through a linear transformation, thereby yielding a complete molecular embedding.

A linear transformation is subsequently applied to the embedding calculated for the molecule, which is formalized as:

where x is the input feature set, *A is* the weight matrix, *b* is the bias, and *y* is the feature vector whose fixed length is set by the user. This transformation standardizes the output length, ensuring uniformity irrespective of the molecule’s size.

The D-MPNN model is trained using the chemprop package (https://github.com/chemprop/chemprop), where parameters are optimized iteratively through forward passes, loss computation, backpropagation, and parameter updates across both the MPNN and a connected feed-forward network (FFN). Our model configuration is guided by the insights [67] enhanced further through Bayesian optimization to refine model performance as detailed in Table 1. Through these settings, our model is able to effectively aggregate and interpret messages from neighbouring atoms, ensuring robust learning across varying chemical structures.

The D-MPNN’s capacity for feature generation and the subsequent training process enable the construction of a predictive model uniquely tailored to the complexity of chemical compounds. By preserving the integrity of molecular graphs and applying nuanced transformations, D-MPNN stands as a powerful tool for advancing our understanding and predictions within chemical informatics.

### 2.2 Performance metrics

To assess the predictive accuracy of our regression model on the test set, we employed the sklearn library to calculate several crucial performance metrics, comparing observed (*y_obs_*) and predicted (*y_pred_*) pGI_50_ values. These metrics are instrumental in quantifying the model’s ability to accurately predict unseen data and provide a comprehensive overview of its overall performance.

Pearson’s correlation coefficient (*Rp*) quantifies the degree of linear relationship between two variables, providing insight into both the strength and direction of their linear association. It is computed as:

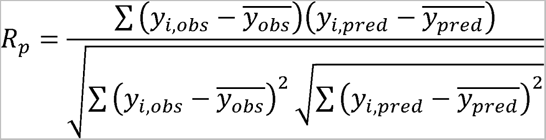

The Rp value varies between −1 and 1. A value approaching 1 indicates a robust positive linear correlation, −1 signifies a strong negative linear relationship, and 0 suggests no linear correlation.

Root Mean Squared Error (RMSE) quantifies the average magnitude of the prediction error, representing how closely a model’s predictions match the actual observed outcomes. It is calculated as the square root of the average squared differences between the predicted and observed values, making it sensitive to large errors. The formula is as follows, where N is the number of observations:

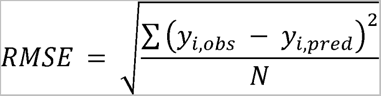

The range of RMSE values is from 0 to infinity. An RMSE value of 0 represents a perfect fit, indicating that predicted values perfectly match the actual values.

Matthews correlation coefficient (MCC) offers a balanced measure for the quality of binary classifications, effectively handling imbalanced datasets. It considers all quadrants of the confusion matrix:

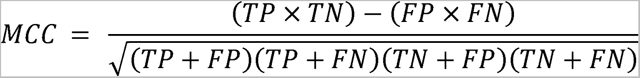

The values of MCC range between −1 and 1, with a score of 1 signifying perfect classification, 0 indicating random classification, and −1 representing complete disagreement between prediction and observation.

Hit Rate (HR), or precision, is the proportion of positive identifications that were actually correct. The HR is calculated as follows and generally expressed in percentages.

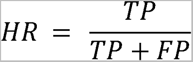

### 2.3 Model validation strategies

Evaluating model performance is paramount in ML, as it ensures a model’s reliability in predicting outcomes on unseen data. Our study utilized two model validation strategies, specifically designed to align with our dataset’s unique characteristics and our research objectives. These strategies aimed to assess the model’s predictive accuracy in diverse scenarios, highlighting its potential for broad application.

#### 2.3.1 Random split

Our initial approach involved a random-split strategy for each cell line within our dataset. In this method, the data for each cell line was divided into two subsets: a training set, which included 90% of the data, and a test set, which comprised the remaining 10%. This partitioning approach served as a foundational step towards implementing a 10-fold cross-validation (CV), facilitating early-stage exploratory analysis and model calibration. By ensuring the random selection of the test set, we aimed to evaluate the model’s performance across a varied sample of the data, thereby promoting a comprehensive understanding of its training effectiveness and predictive accuracy.

#### 2.3.2 Dissimilar-molecules split

To enhance the rigor of our model evaluation, we implemented a strategy focusing on the inclusion of dissimilar molecules during the dataset partitioning stage [18], [40]. After performing an initial random division of the dataset, we analysed the structural similarity of the molecules using SMILES notation to differentiate those in the training and test sets. We then applied a similarity threshold: molecules within the test set showing more than 70% similarity to any molecule in the training set were removed. This approach aimed to guarantee that the test set was composed solely of molecules significantly distinct from those the model was trained on. By adopting this methodology, we sought to rigorously assess the model’s ability to generalize its predictions to new, chemically unrelated compounds, thereby validating its effectiveness in identifying and assessing novel compounds.

## 3 Results

### 3.1 Comparative performance of ML algorithms on unseen randomly split NCI-60 dataset

In this study, we evaluated a random-split model to predict the pGI50 responses across various cancer cell lines for a comprehensive set of compounds, utilizing the extensive molecular screening data available from the NCI-60 dataset. While previous studies [22], [23], [34], [39], [42] have focused on smaller datasets, our research utilizes a much larger dataset, which includes data across 60 cell lines corresponding to nine cancer types. This extensive data volume provides a robust foundation for our ML models, which is pivotal for achieving superior predictive performance in phenotypic virtual screening. Employing algorithms such as LR, RF, XGB, DNN, and D-MPNN, we trained models for each cell line to predict pGI_50_ based on compounds features.

The ML models underwent training using the recommended values for their hyperparameters shown in Section 2.1, followed by a hyperparameter-tuning phase to evaluate potential performance enhancements (see Table 1 for reference). The LR algorithm was trained exclusively with the recommended hyperparameters. In Figure 3, we compare the results of both scenarios, where each algorithm’s performance is depicted in a boxplot that includes 60 scores (one per cell line) of either RMSE or Rp.

**Figure 3.**
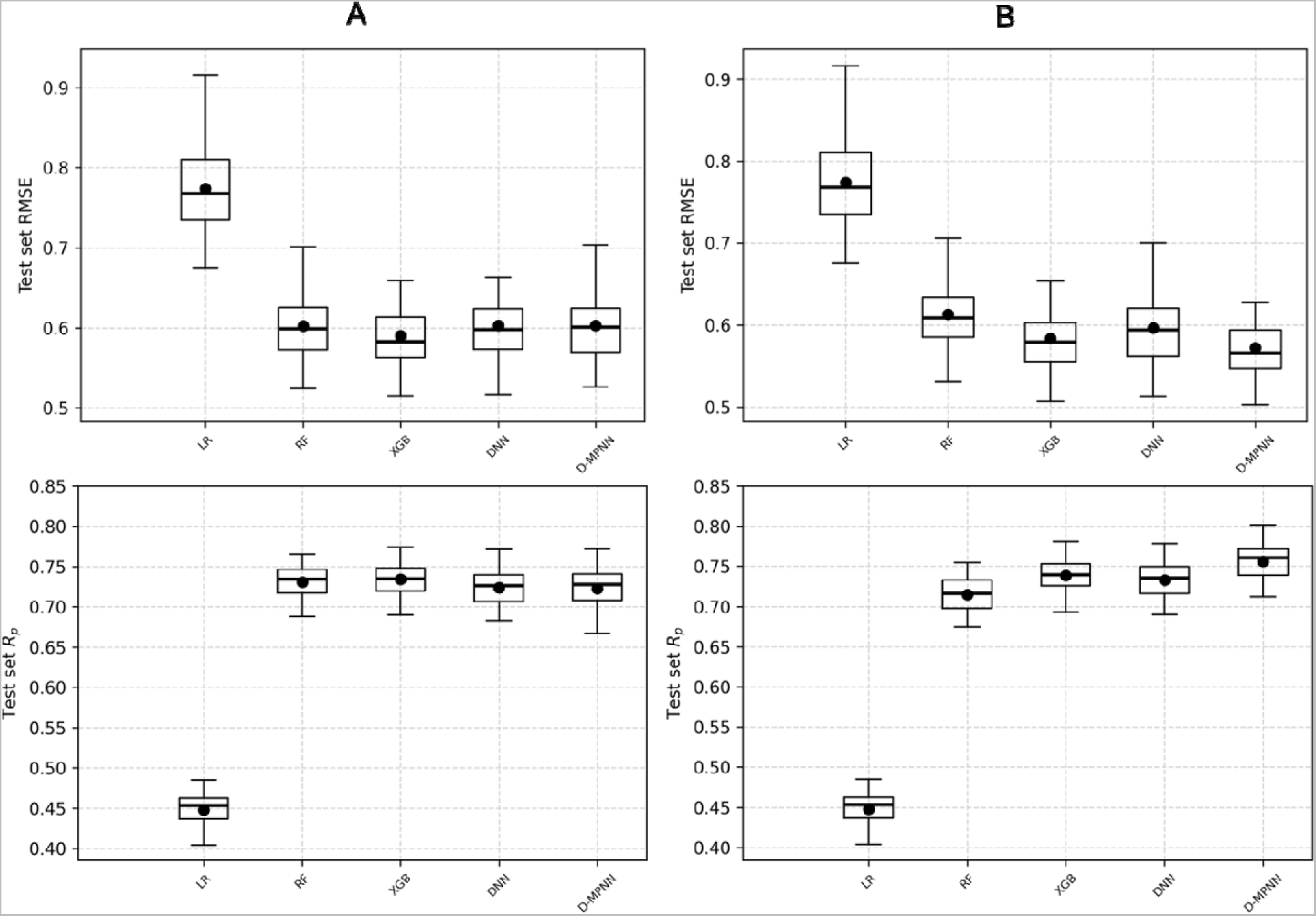
Hyperparameter tuning generally resulted in better performance. The performance of each algorithm across NCI-60 cell lines was evaluated using a random split. Results were obtained using (A) recommended hyperparameters and (B) hyperparameters acquired after tuning. Each subpanel shows the results for the following ML algorithms from left to right: LR, RF, XGB, DNN, and D-MPNN. The boxplot for each algorithm contains 60 scores, either RMSE (upper panel) or Rp (lower panel). The central line of each boxplot represents the median, and the dot denotes the mean. D-MPNN benefitted the most from hyperparameter tuning.

With the recommended values for its hyperparameters, the XGB models outperformed the other four model types in both RMSE and Rp metrics. Hyperparameter tuning notably enhanced the performance of the XGB models, highlighting the significance of hyperparameter interactions in gradient boosting methods. However, the D-MPNN models outperformed the XGB models after hyperparameter tuning, with the XGB models coming in second. Hyperparameter tuning had a minimal impact on the performance of RF models, consistent with prior literature suggesting their insensitiveness to hyperparameter tuning [51]. Using recommended hyperparameter values, the baseline LR models again exhibited much lower accuracy compared to its nonlinear counterparts.

The DNN models also obtained improved performance upon hyperparameter tuning. The analysis revealed that a constant learning rate and an increased number of neurons per hidden layer contribute to this enhancement. These findings challenge previous recommendations [64] and highlight the importance of dataset-specific hyperparameter tuning.

After hyperparameter tuning, the D-MPNN models achieved the most substantial improvement, reaching the lowest average RMSE of 0.572 and the highest average Rp of 0.756. Thus, this shows that D-MPNN is the best algorithm for these datasets. Bayesian optimization [68] applied to D-MPNN for hyperparameter tuning due to its extensive hyperparameter space yielded significant improvements. However, for other algorithms, Bayesian optimization did not offer advantages over grid search (data not shown).

In conclusion, our comprehensive assessment indicates that while XGB models are efficient and perform well, D-MPNN models, when optimally tuned, are the most proficient in capturing the complex relationships in the data, as demonstrated by their superior RMSE and *R_p_* scores.

### 3.2 Models performance on dissimilar molecules

We extended our investigation into model robustness by evaluating their performance on a dissimilar test set, defined by a 70% dissimilarity threshold. This critical benchmark was designed to emulate the challenge of predicting the efficacy of novel compounds and to validate the generalizability of our models to chemical entities vastly different from those seen during training. The dissimilarity criterion is pivotal in ensuring that our models are not merely reflecting the chemical space of the training set. We meticulously evaluated the predictive accuracy and error margins of each ML algorithm on this challenging set. Following the initial results, the models were trained in this phase using their optimized hyperparameters.

The findings in Figure 4 align with those in Figure 3, indicating that the linear regression model consistently underperforms compared to its nonlinear counterparts. Notably, the D-MPNN model demonstrates the best results, with the highest mean Rp of 0.632 and the lowest mean RMSE of 0.584, highlighting its strong generalization capabilities.

**Figure 4.**
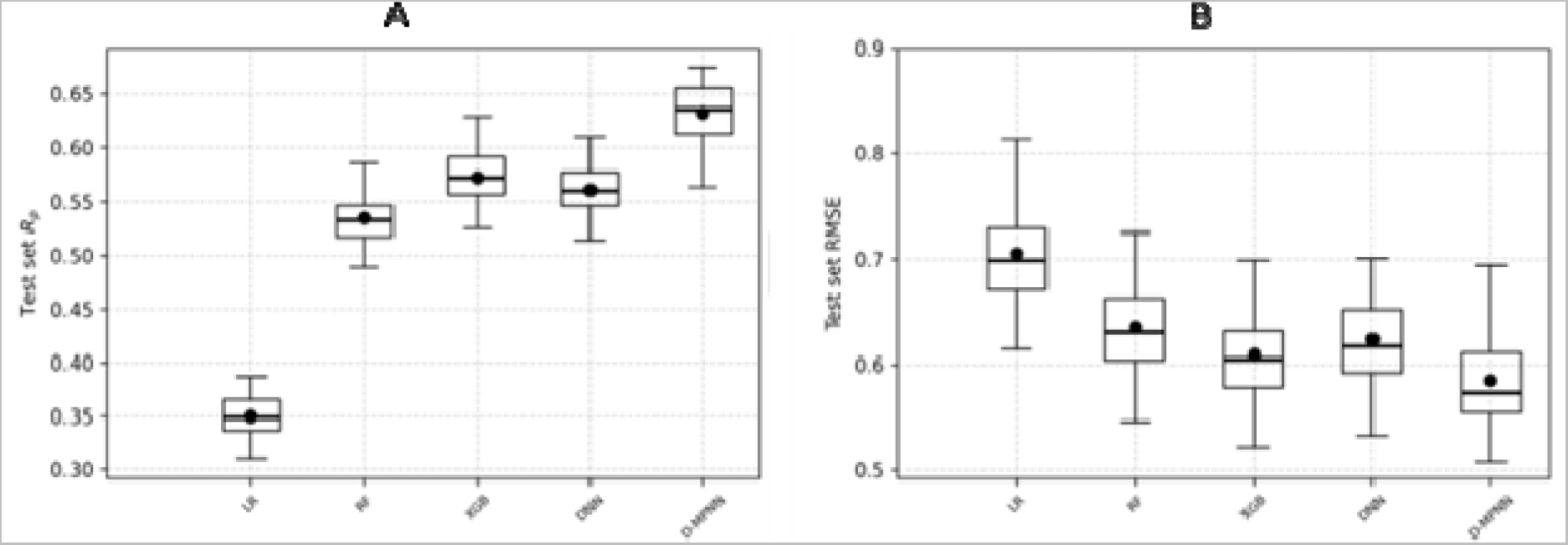
D-MPNN also performs best on the dissimilar-molecules test sets. The performance of each algorithm, using their optimized hyperparameters, across NCI-60 cell lines was evaluated on the dissimilar test set. Each subpanel shows the results for the following ML algorithms from left to right: LR, RF, XGB, DNN, and D-MPNN. The boxplot for each algorithm contains 60 scores, either (A) Rp or (B) RMSE. The central line of each boxplot represents the median, and the dot denotes the mean.

To emphasize the importance of implementing a robust model validation strategy, especially in the early stages of drug discovery, we summarized the performance of the models across 60 cell lines by averaging the RMSE and Rp scores. Note that these values correspond to the dot markers shown in Figures 3 and 4. The results of this comparison are presented in Table 3, showing that models perform better when using random data partitioning compared to partitioning based on dissimilar molecules. The reason for this performance reduction is that in the first case, the test set may contain molecules similar to those in the training set, which allows the models to recognize these molecules. Conversely, in the second case, partitioning based on dissimilar molecules ensures that the test set only contains molecules that are different from those in the training set, making it a more challenging scenario and thus reducing the performance of the ML models.

**Table 3.** Comparison of the two model validation strategies. In both, random split and dissimilar-molecules split. The performance of each ML model is summarized using the average of its RMSE or Rp scores. Results presented here correspond to models trained with optimized hyperparameters, except for the LR model.

| ML algorithms | Random split |  | Dissimilar split |  |
| --- | --- | --- | --- | --- |
| | $R_p$ | RMSE | $R_p$ | RMSE |
| LR | 0.4480 | 0.7742 | 0.3499 | 0.7052 |
| RF | 0.7150 | 0.6131 | 0.5344 | 0.6356 |
| XGB | 0.7385 | 0.5838 | 0.5727 | 0.6093 |
| DNN | 0.7331 | 0.5963 | 0.5613 | 0.6242 |
| D-MPNN | **0.7563** | **0.5724** | **0.6326** | **0.5850** |

Despite its reduced accuracy on dissimilar-molecules split, the D-MPNN model outperforms all other ML models. Indeed, D-MPNN’s pGI_50_ predictions have lower error and higher correlation when compared with the corresponding measurements across both validation strategies, highlighting its potential to drive early drug design approaches.

These results underscore the critical importance of robust and stringent testing environments for the validation of ML models within the field of drug discovery. The standout predictive performance of the D-MPNN model on the dissimilar test set underscores its ability to effectively generalize to novel chemical entities, a key attribute for models applied to the prediction of compound efficacy. The D-MPNN’s superior performance in such complex predictive scenarios confirms its status as a potent tool in the arsenal of computational drug discovery.

### 3.3 D-MPNN model performance in phenotypic virtual screening

Figure 5 provides a compelling demonstration of the D-MPNN model’s strength in the context of phenotypic virtual screening, particularly in its ability to provide a high hit rate across both explored (random split) and unexplored chemical spaces (dissimilar-molecules split). The figure illustrates predictive performance on the best and worst cell lines as determined by the pGI_50_ cut-off, used to differentiate between inactive (pGI_50_ < 6) and active (pGI_50_ ≥ 6) compounds.

**Figure 5.**
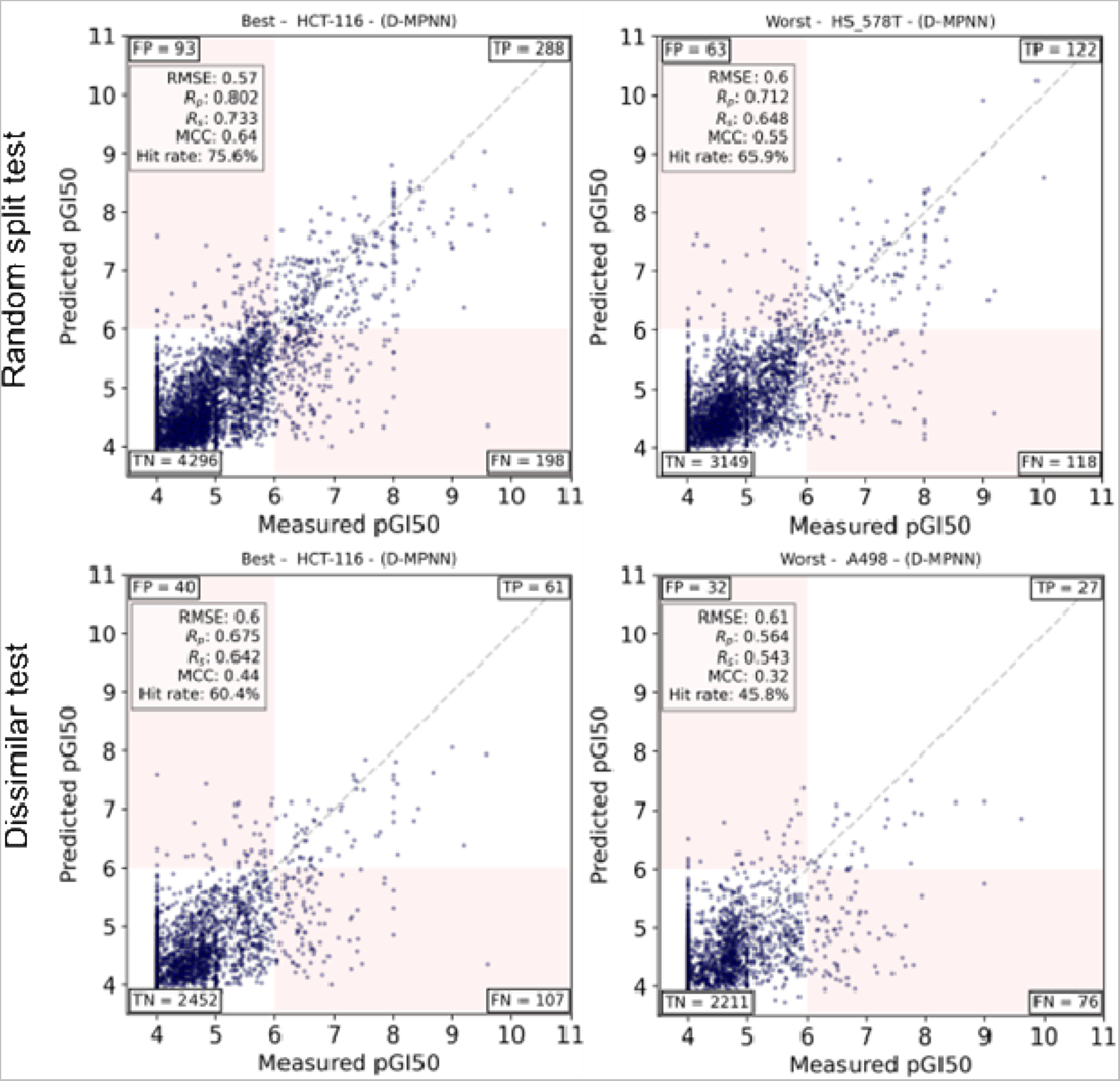
D-MPNN model performance in phenotypic virtual screening using either the random or the dissimilar-molecules split. For each cell line, the same training set is employed regardless of the split, whereas the test set either contains only those dissimilar to those in the training set or all the remaining molecules (both dissimilar or similar). Both regression and classification performance is simultaneously evaluated using a pGI50 cutoff of 6 for the latter. The scatter plots compare predicted pGI50 values against their measured values for the RMSE-best and RMSE-worst cell lines (left and right, respectively) under random split (top panels) and dissimilar-molecules split (bottom panels). Notably, the model maintains high hit rate, Rp and MCC values with the dissimilar-molecules split. These results highlight D-MPNN’s robustness and its capacity to identify potent molecules with potent whole-cell inhibition among a much larger proportion of inactive molecules, thus demonstrating its value for virtual screening against cancer cell lines.

In the explored setting of the random split test, the D-MPNN showcases its high predictive fidelity on the best cell line, HCT-116, achieving a *R_p_* score of 0.802, which is indicative of a strong linear correlation between predicted and measured values. The high hit rate of 75.6% and an MCC of 0.64 in this scenario underscore the model’s precise ability to identify potent compounds—a critical requirement for effective virtual screening.

The robust nature of the D-MPNN model is further emphasized in the dissimilar test set, where even the best cell line, HCT-116, presents a new challenge due to the introduction of dissimilar molecules. Despite this, the model attains a *R_p_* score of 0.675, a substantial hit rate of 60.4%, and an MCC of 0.44. These metrics are particularly good, considering the complexity posed by the dissimilarity threshold, and they highlight the D-MPNN’s adaptability and its capacity to recognize active agents in a less explored chemical space.

The worst-performing cell line in the dissimilar test, A498, while exhibiting lower metrics *R_p_* of 0.564 and hit rate of 45.8%—still demonstrates the D-MPNN model’s relative resilience. An MCC of 0.32 in this context demonstrates the model’s ability to maintain a fair level of true positive identification amidst increased chemical diversity, which is often encountered in real-world screening environments.

## 4 Discussion

Our study aimed to identify the most effective AI-based model for predicting the inhibitory activity of molecules, especially molecules that differ from those encountered during the training process. The models were trained using the NCI-60 dataset. In this dataset, cell lines are tested with at least 30,000 molecules (see Figure 1), which allows for a robust assessment of model performance across different cancer cell lines. This approach differs from many studies that use much smaller and less diverse datasets, which may limit their applicability and generalizability [35], [70]. This study suggests that larger, more heterogeneous datasets can significantly enhance the model’s ability to generalize, supporting previous findings that dataset diversity is crucial for training effective AI models in drug discovery applications [11], [12], [15].

When comparing the two data partitioning strategies for model training and testing, we find that models tend to perform better when we use random data partitioning instead of partitioning based on dissimilar molecules (see Table 3). This is because when we use random data partitioning, the test set also includes molecules similar to those in the training set, allowing the models to predict those more accurately. On the other hand, only using dissimilar molecules in the test set makes the task more challenging for the models, leading to decreased performance in ML models.

Regardless of the strategy used, our results showed that hyperparameter tuning improves the performance of all the models evaluated in this study (Figures 3 and 4). Moreover, this study has demonstrated that D-MPNNs offer superior predictive capabilities in virtual screening for anticancer therapeutics against diverse cancer cell lines, even when analysing chemically dissimilar molecules. Our findings align with recent advancements in the field that suggest GNNs’ superior ability to model complex molecular interactions due to their inherent structure that mimics molecular graphs [34], [69]. In contrast with other ML models like RF and XGB, which do not inherently capture the topological variability of molecular structures as effectively. Taken together, these results suggest the usefulness of D-MPNN for phenotypic virtual screening applications. The high hit rates and MCC scores show the D-MPNN’s inherent strength in uncovering compounds with strong inhibitory activity against cancer cell lines, thereby supporting its use in the crucial early stages of drug discovery where the prediction of compound efficacy is paramount.

## 5 Conclusions

The results of this study demonstrate the significant role of GNNs as a powerful tool in virtual screening technologies, which aim to improve the discovery of anticancer drugs. GNNs are able to accurately predict the biological activity of chemically diverse molecules against different cancer cell lines, demonstrating a capability to navigate complex chemical spaces that outperform that of other types of AI models struggle to explore fully.

We conducted evaluations using the NCI-60 dataset, one of the largest and most diverse collections in such studies. Our findings suggest that GNNs are robust and highlight the crucial role of using comprehensive datasets to enhance the accuracy and generalizability of predictive models in oncology. This research contributes to our understanding of how various AI models can be optimized and tailored to address the specific requirements of early drug design, thus expanding the possibilities of what can be achieved with current technologies.

Future research should concentrate on refining specialized models, utilizing transfer learning to capitalize on knowledge derived from various cancer types. This approach can potentially improve predictive accuracy by allowing models to generalize across diverse cancer datasets. Additionally, the integration of multi-omics data with chemical data via multi-task learning could substantially improve virtual screening performance against cancer cell lines. Evaluation of other GNNs on the 60 dissimilar-molecules splits across NCI-60 cell lines is also promising.

Code for reproducing the results of the best models can be found at https://github.com/sachin-vish91/GNN-VS

## Acknowledgments

S.V. thanks the Institute Paoli Calmettes Marseille, France for his PhD funding. S.H-H. thanks the National Council of Sciences and Technology of Mexico (CONAHCYT). P.J.B. thanks the Wolfson Foundation and the Royal Society for a Royal Society Wolfson Fellowship. Part of the computational experiments were performed on the Core Cluster of the French Institute of Bioinformatics (IFB) (ANF-11-INBS-0013), which is also gratefully acknowledged.

